# Groucho/TLE opposes axial to hypaxial motor neuron development

**DOI:** 10.1101/2020.03.10.986323

**Authors:** Adèle Salin-Cantegrel, Rola Dali, Jae Woong Wang, Marielle Beaulieu, Mira Deshmukh, Ye Man Tang, Stefano Stifani

## Abstract

Spinal cord motor neuron diversity and the ensuing variety of motor circuits allow for the processing of elaborate muscular behaviours such as body posture and breathing. Little is known, however, about the molecular mechanisms behind the specification of axial and hypaxial motor neurons controlling postural and respiratory functions respectively. Here we show that the Groucho/TLE (TLE) transcriptional corepressor is a multi-step regulator of axial and hypaxial motor neuron diversification in the developing spinal cord. TLE first promotes axial motor neuron specification at the expense of hypaxial identity by cooperating with non-canonical WNT5A signalling within the motor neuron progenitor domain. TLE further acts during post-mitotic motor neuron diversification to promote axial motor neuron topology and axonal connectivity whilst suppressing hypaxial traits. These findings provide evidence for essential and sequential roles of TLE in the spatial and temporal coordination of events regulating the development of motor neurons influencing posture and controlling respiration.

**HIGHLIGHTS:**

- Groucho/TLE mediates non-canonical WNT signalling in developing motor neurons
- Non canonical WNT:TLE pathway regulates thoracic motor neuron diversification
- TLE promotes axial while inhibiting hypaxial motor neuron development
- TLE influences developing motor neuron topology and muscle innervation

**IN BRIEF:** Salin-Cantegrel et al use *in ovo* engineered approaches to show that a non-canonical WNT:TLE pathway coordinates temporally and spatially separated elements of motor neuron diversification, repressing hypaxial motor neuron development to promote the axial fate.

**GRAPHICAL ABSTRACT:** TLE contribution to the development of thoracic somatic motor columns
Progenitor cells in the ventral pMN domain are exposed to higher concentrations of non-canonical WNTs and express more TLE. Cooperation of non-canonical WNTs and TLE renders ventral pMN progenitors refractory to a respiratory MN fate, thereby contributing to the separation of MMC and RMC MN lineages. Differentiating MNs that maintain high TLE expression also maintain LHX3 expression, adopt axial motor neuron topology and connect to axial muscles. TLE activity in differentiating MMC MNs prevents the acquisition of respiratory MN topology and innervation traits.

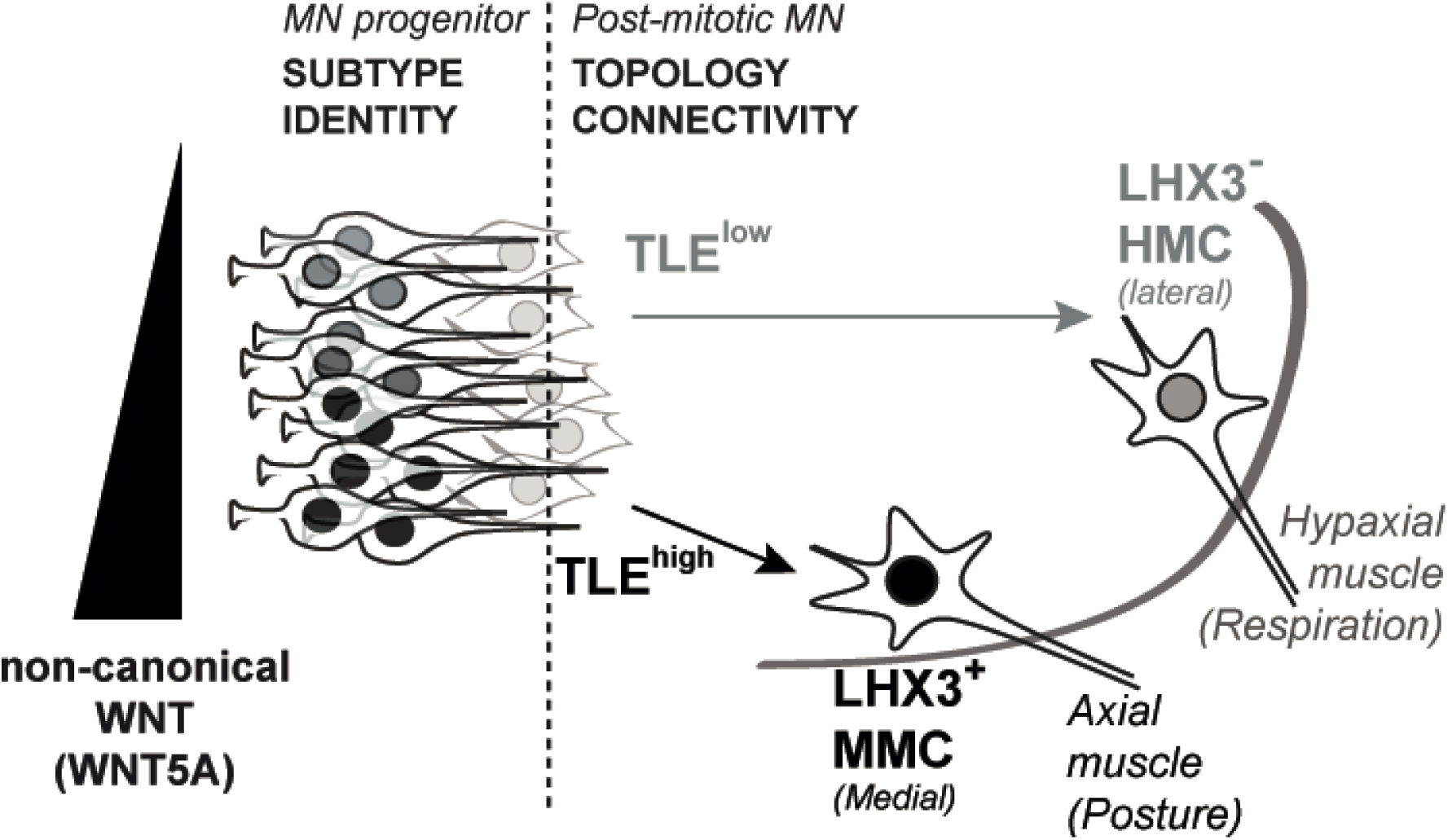

## INTRODUCTION

Axial and hypaxial muscles, which control undulatory locomotor behaviour in primitive chordates, have evolved in more complex organisms to sustain erectile body posture or control vital functions such as respiration. As a result, vertebrates from birds to mammals share the feature of using axial muscles to maintain their stance while hypaxial muscles assist breathing ^1,2^. Axial muscle-innervating motor neurons (MNs) are usually found along the entire spinal cord length, whereas phrenic and hypaxial muscle-innervating MNs in mammals, are segmentally restricted to cervical and thoracic levels ^3-6^. Although most breathing functions are controlled by phrenic MNs and the diaphragm in human, a quarter of respiration depends primarily on hypaxial MNs capacity to contract intercostal muscles, allowing the thoracic cavity and the lungs to expand and deflate. Thus, selective death of hypaxial MNs and their defective function in disease cause devastating ventilatory insufficiencies^7^. Despite their vital biological role, little is presently known about the mechanisms controlling the diversification of hypaxial respiratory MNs during spinal cord development.

MN development is a multi-step process that takes place both prior to, and following, acquisition of post-mitotic neuronal features. Within the ventral germinative zone of the neural tube, the transcription factors NKX6.1, PAX6 and OLIG2 are critical to delineate the boundaries of the MN progenitor (pMN) domain ^8-12^ and to regulate the expression of transcription factors HB9, LHX3 and ISL1, whose combined activity is necessary for the implementation of a generic MN developmental program ^13-15^. At thoracic spinal cord level, axial MNs of the median motor column (MMC), which maintain sustained LHX3 expression during their maturation, are specified by a ‘ventral^high^ to dorsal ^low^’ gradient of non-canonical WNT molecules, such as WNT4, WNT5A and WNT5B ^16^. In contrast, MNs of the hypaxial motor column (HMC) extinguish LHX3 expression during development and are thought to be derived from progenitor cells refractory to graded non-canonical WNT signalling and responsive to competing dorsal canonical WNT activity ^16,17^. The precise molecular mechanisms underlying the roles of WNT signalling pathways in the separation of MMC and HMC MN fates are, as of yet, poorly characterized.

Canonical WNTs bind to specific cell surface receptor/co-receptor complexes allowing the interaction of β-CATENIN with members of the TCF/LEF (TCF hereafter) family of HMG-box DNA-binding transcription factors, leading to β-CATENIN:TCF-mediated transactivation of target genes ^18^. In contrast, non-canonical WNT pathways are β-CATENIN-independent and have distinct or even opposite functions to canonical WNT signalling ^19-22^. It has been proposed that non-canonical WNTs, such as WNT5A, can antagonize β-CATENIN:TCF-mediated transactivation and instead promote the functions of TCF-containing transcriptional repressor complexes ^23,24^. In this regard, important TCF cofactors with transcription repression activity include members of the evolutionarily conserved Groucho/TLE (TLE hereafter) co-repressor family ^25-28^.

Here we provide evidence that TLE antagonizes canonical, and promotes non-canonical, WNT signalling in the developing spinal cord. TLE and non-canonical WNT signalling work together in MN progenitors to promote MMC diversification at the expense of the HMC fate. Furthermore, TLE also acts in post-mitotic MNs to further specify MMC neuronal settling position and axonal connectivity at the expense of HMC topology and innervation. These findings provide information about the mechanisms regulating the axial versus hypaxial MN binary fate choice, offering new insight into the molecular programs governing the generation of MNs controlling posture and respiration.

## RESULTS

### TLE modulates WNT pathways in the developing thoracic spinal cord

Previous studies have shown that TLE is important for ventral spinal tube patterning and is expressed in MN progenitors at developmental stages coinciding with spinal MN specification ^9,29^. To begin to study the involvement of TLE in MN development, we first characterized its expression in the developing chick ventral spinal cord at stages following neural tube patterning. Using a previously described ‘panTLE’ monoclonal antibody recognizing all members of the TLE protein family ^29-31^, we observed TLE^+^ cells in the progenitor zone in the ventral thoracic spinal cord of Hamburger and Hamilton (HH) stage 21 chick embryos (Figure 1A and 1B); a dynamic ventra ^high^-to-dorsal^low^ TLE immunoreactivity was found across the NKX6.1^+^ progenitor domains, including the OLIG2^+^ pMN domain (Figure 1A and Figure S1). Except for the most ventral cells, the majority of TLE^+^ progenitors robustly coexpressed the WNT pathway transcription factor TCF4, a member of the TCF/LEF family of TLE binding proteins ^25,32,33^ (Figure 1B). These observations prompted us to assess whether TLE might be involved in modulating WNT activities in the developing spinal cord.

**FIGURE 1.**
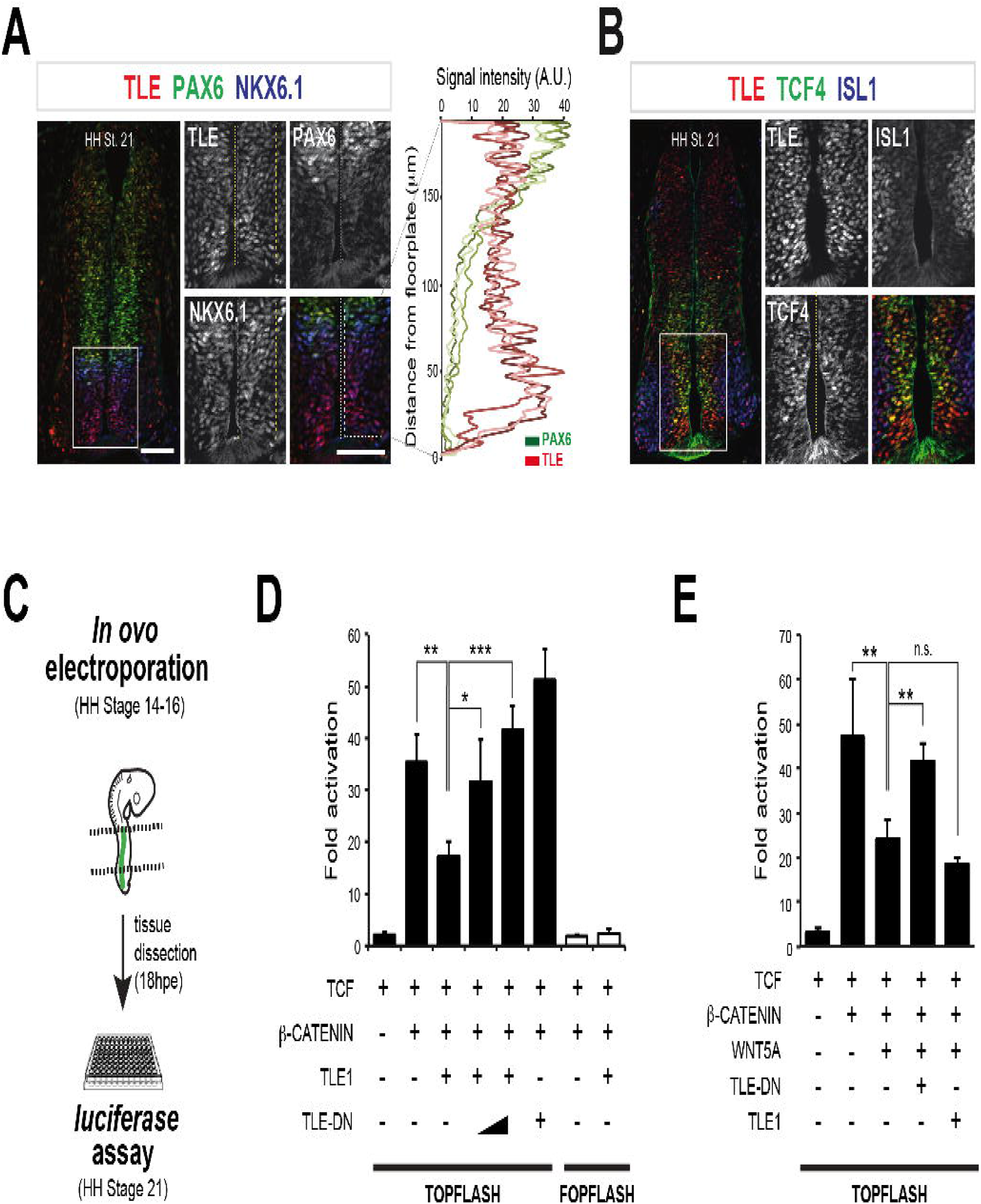
Non-canonical WNT and TLE functional interaction in the developing spinal cord. (A) Expression of TLE, PAX6, and NKX6.1 in HH stage 21 chick embryonic spinal cord. Also shown is fluorescence signal intensity of TLE and PAX6 protein levels quantified within both NKX6.1+ ventral progenitor regions of HH Stage 21 chick embryos (n=3 embryos). The *x*-axis depicts signal level of TLE and PAX6 and the *y*-axis the distance (μm) from the floor plate. Here and in all succeeding figures, boxed areas are shown at higher magnification in columns depicting single channels and scale bars correspond to 100 μm. (B) Expression of TLE, TCF4, and ISL1 in HH stage 21 chick embryonic spinal cord. (C) Schematic representation of *in ovo* transcription assays used in the following experiments. (D and E) Quantification of luciferase activity in chick embryonic spinal cord cells electroporated with TOPFLASH or FOPFLASH (negative control) *luciferase* reporter constructs and plasmids expressing the indicated combinations of proteins. Results are shown as mean ± SEM (n≥4 sets of experiments, \**p* < 0.05; \*\**p* < 0.01; \*\*\**p* < 0.001; n.s., not significant; two-way ANOVA).

To begin to examine this possibility, chick embryo thoracic spinal cords were *in ovo* electroporated at HH stage 14-16 with a previously described ^32^ β-CATENIN:TCF-responsive reporter construct driving *luciferase* gene expression from tandem TCF/LEF binding sites in order to monitor canonical WNT activity (Figure 1C). Coexpression of β-CATENIN and TCF resulted in robust *luciferase* transactivation in spinal cord cells (Figure 1D, bars 1 and 2). Expression of full-length TLE1 (‘CMV∷TLE1’) significantly reduced β-CATENIN:TCF-mediated transactivation (Figure 1D, bars 2 and 3). This inhibitory effect of TLE1 was blocked in a dose-dependent manner by a previously characterized ^29^ TLE1 deletion mutant that does not have transcriptional co-repressor ability, thereby acting as a dominant-negative (‘DN’) form (‘TLE-DN’ hereafter) (CMV∷TLE1+CMV∷TLE-DN, *p* = 0.00012 compared to CMV∷TLE1 alone, at highest CMV∷TLE-DN amount tested; ANOVA followed by Bonferroni *post hoc* test; n≥4 embryos) (Figure 1D, bars 3-5). A trend toward increased transactivation was observed when TLE-DN was expressed in the absence of TLE1 (Figure 1D, bar 6). No detectable transcriptional effects were noted when a mutated, negative control, reporter construct was used (Figure 1D, bars 7 and 8). These results suggest that TLE can antagonize canonical WNT in the developing spinal cord. In agreement with this possibility, we noticed an almost inverse correlation between the expression of TLE and PAX6, a transcriptional target of β-CATENIN:TCF-mediated transactivation ^34^ in the ventral spinal cord region (Figure 1A).

We then examined whether TLE might be involved in the modulation of non-canonical WNT signalling in the developing spinal cord. To do so, we determined the impact of endogenous TLE on the ability of WNT5A to antagonize β-CATENIN:TCF-mediated transactivation *in vivo*. Exogenous expression of WNT5A (‘CMV∷WNT5A’) significantly reduced β-CATENIN:TCF-mediated transactivation (Figure 1E, bars 1-3) and this effect was blocked by coexpression of TLE-DN (CMV∷WNT5A+CMV∷TLE-DN, *p* = 0.0017 compared to CMV∷WNT5A alone; ANOVA followed by Bonferroni *post hoc* test; n≥4 embryos) (Figure 1E, bars 3 and 4). CMV∷WNT5A *in ovo-*electroporation also resulted in a detectable decrease in PAX6 immunoreactivity in the ventral PAX6^+^ domain (Figure S2A and S2B), mimicking the suppressive effect of exogenous TLE on ventral PAX6 expression (Figure S2C, and as previously shown^29^). Together, these data suggest that TLE has the ability to antagonize canonical, and promote non-canonical, WNT pathways in the developing spinal cord.

### TLE in non-canonical WNT-mediated specification of axial as opposed to hypaxial motor neurons

We next investigated whether TLE was involved in the non-canonical WNT ability to regulate somatic MN subtype specification^16^. HH stage 14-16 chick thoracic spinal cords were *in ovo* electroporated with green fluorescent protein (GFP) alone or together with CMV∷WNT5A in the absence or presence of CMV∷TLE-DN, followed by quantification of the numbers of somatic (HB9 and ISL1 co-expression) MNs exhibiting either a MMC (HB9^+^/ISL1^+^/LHX3^+^) or HMC (HB9^+^/ISL1^+^/LHX3^-^) molecular profile at thoracic level (Figure 2A-C). Exogenous expression of WNT5A caused a significant increase in neurons with a MMC molecular profile 96 hours post electroporation (hpe) (HH stage 30) (*p* = 0.0082, ANOVA followed by Tukey *post hoc* test; n=5-6 embryos), with a parallel decrease in neurons with a HMC profile (*p* = 0.0114, ANOVA followed by Tukey *post hoc* test; n=5-6 embryos) (Figure 2B, bars 1 and 2, 4 and 5). These WNT5A-mediated effects were blocked when endogenous TLE function was inhibited by TLE-DN coexpression (*p* = 0.026 compared to WNT5A embryos; ANOVA followed by Tukey *post hoc* test; n=5-6 embryos) (Figure 2B, bars 2 and 3, 5 and 6). No detectable consequences on either the numbers of visceral MNs of the *Column of Terni* (ISL1^+^ but negative for both HB9 and LHX3) (Figure 2B, bars 7-9) or total MNs (Figure 2C) were observed. We also looked closer at whether TLE-DN might be sufficient to antagonize endogenous non-canonical WNT signalling and its *in vivo* effect of MN subtypes specification through *in ovo* electroporation of CMV∷TLE-DN alone. To do so, MMC MNs were quantified on the electroporated side of the spinal cord and compared to the number of their non-electroporated controlateral (control) counterparts at 96 hpe (Figure 2D). The expression of TLE-DN (Figure 2E) elicited a decrease in MNs with a MMC identify (*p =* 0.043 compared to control side, Student’s *t* test; n=7 embryos). This change in MMC MN number was paralleled by an increase in MNs with a HMC identity (*p* = 0.00016 compared to control side, Student’s *t* test; n=7 embryos) (Figure 2E, bars 1-4). No detectable effects on visceral MN numbers were observed (Figure 2E, bars 5 and 6). A modest decrease in total MN number was noticed after TLE-DN electroporation (Figure 2F). Thus, TLE may possibly act downstream of non-canonical WNT signalling during the specification of the MMC, as opposed to HMC, fate in the developing chick thoracic spinal cord.

**FIGURE 2.**
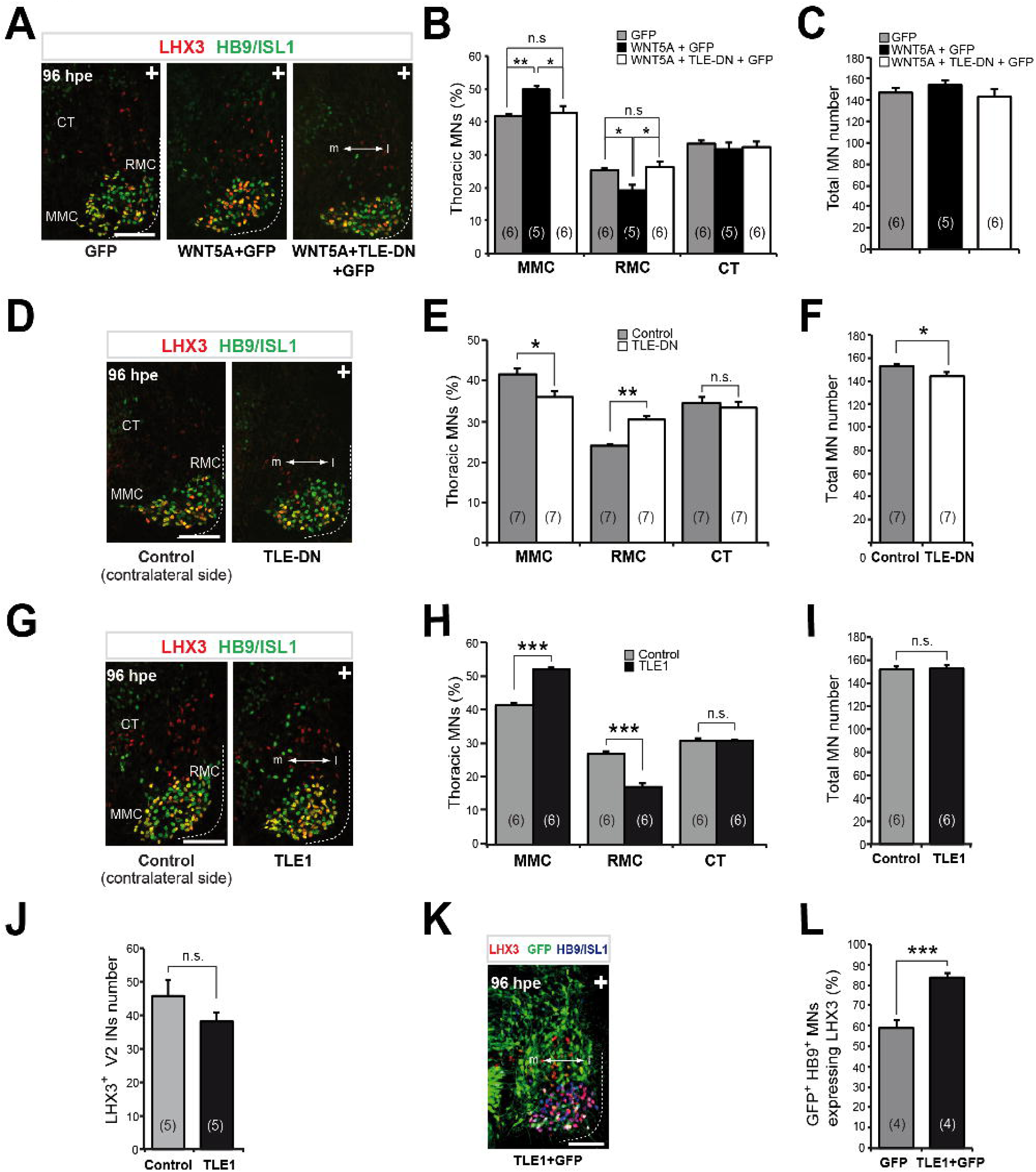
Involvement of TLE in non-canonical WNT signalling during development of thoracic motor neurons. (A) Expression of LHX3, HB9/ISL1 and GFP in representative hemi-sections of chick thoracic spinal cords subjected to *in ovo* electroporation with plasmids expressing GFP alone, WNT5A+GFP, or WNT5A+TLE-DN+GFP as indicated and then analysed 96 hpe (HH Stage 30). Here and in succeeding figures “m” and “l” stand for medial and lateral sides of the spinal cord and “+” indicates the electroporated side. (B and C) Quantification of somatic thoracic MN columnar subtypes (B) and total thoracic MN numbers (C) under the indicated experimental electroporation conditions (here controls correspond to proportions observed in GFP-electroporated embryos). Results are shown as mean ± SEM (\**p* < 0.05; \*\**p* < 0.01; n.s., not significant; two-way ANOVA). Here and in succeeding figures the number of embryos analyzed is indicated in brackets. (D) LHX3 and HB9/ISL1 expression in hemi-sections of chick spinal cords subjected to *in ovo* electroporation with CMV:TLE-DN and then analysed 96hpe. (E and F) Proportion of thoracic MN columnar subtypes (F) and total MN numbers (E) per ventral quadrant in TLE-DN electroporated embryos compared to the non-electroporated side (control). Results are shown as mean ± SEM (\**p*<0.05; \*\**p*<0.01; n.s., not significant; *t* test). (G) LHX3 and HB9/ISL1 expression in chick embryos spinal cord *in ovo* electroporated with CMV∷TLE1 and analysed 96 hpe. (H-J) Quantification of the proportion of thoracic motor neuron columnar subtypes (H), total MN numbers (I), or LHX3+ V2A interneuron numbers (J) per ventral quadrant of TLE1-electroporated embryos compared to the non-electroporated side (control). (K) Expression of LHX3, HB9/ISL1 and GFP, at 96 hpe, in representative spinal cord hemi-sections of chick embryos co-electroporated with TLE1 and GFP. (L) Quantification of thoracic LHX3+ MNs in chick embryo spinal cords electroporated with GFP alone or together with TLE1. Results are shown as mean ± standard errors of the mean (SEM) (\*\*\**p* < 0.001; *t* test).

To examine this possibility further, we tested whether forced exogenous TLE expression would phenocopy the promotion of MMC neuron specification elicited by WNT5A. *In ovo* electroporation-mediated expression of CMV∷TLE1 resulted in an increased number of electroporated somatic MMC MNs (*p* = 0.0000004, Student’s *t* test; n=6 embryos) (Figure 2G, bars 1 and 2). Concomitantly, the number of MNs with a HMC molecular identity decreased on the electroporated half compared to the control half (*p* = 0.0000014, Student’s *t* test; n=6 embryos) (Figure 2H, bars 3 and 4). These phenotypes appeared to be specific since exogenous TLE1 did not alter the proportion of visceral MNs (Figure 2H, bars 5 and 6) and did not significantly change the total number of MNs (Figure 2I) or LHX3^+^/HB9^-^ V2A interneurons (Figure 2J). Furthermore, TLE1 exogenous expression increased the number of GFP co-electroporated somatic MNs with a MMC molecular profile as early as 48 hpe (Figure S3) and at 96 hpe (*p* = 0.00068, Student’s *t* test; n=4 embryos) (Figure 2K and 2L) compared to GFP alone. Finally, exogenous expression of point mutants TLE1(L743F) and TLE1(R534A), which both retain the ability to antagonize canonical WNT pathway but specifically fail to interact with other key TLE partners ^35^, phenocopied wild-type TLE1 exogenous expression, in agreement with a functional interaction of TLE with WNT pathways in MMC neurons specification (Figure S4). Taken together, these findings provide evidence suggesting a role for a non-canonical WNT:TLE pathway in the regulation of axial as opposed to hypaxial MN specification in the chick thoracic spinal cord.

### TLE expression in spinal axial and hypaxial motor neurons

A role for TLE in MN development was then considered by following its expression in postmitotic MNs of HH stages 21-28 chick embryos: these stages encompass subtype diversification, target muscle connection and body settlement of ventral MNs ^3,5,36^. TLE expression was detected in most mantle zone somatic MNs expressing HB9 and ISL1. Some HB9/ISL1-expressing MNs exhibited robust TLE expression [hereafter designated ‘TLE^high^’] (Figure 3A, arrow), while others displayed weaker TLE immunoreactivity [‘TLE^low^’] (Figure 3A and 3B, arrowhead) as early as HH stage 21. TLE^high^ MNs expressed the marker LHX3, suggesting that they are part of the MMC (Figure 3A and 3B). TLE^low^ MNs coexpressing HB9 and ISL1 were found mainly in lateral positions, expressed little or no LHX3 and were retrogradely labelled after injection of cholera toxin B (CT-B) into intercostal muscles, targets of thoracic hypaxial MNs, suggesting that they are part of the HMC (Figure 3C).

**FIGURE 3.**
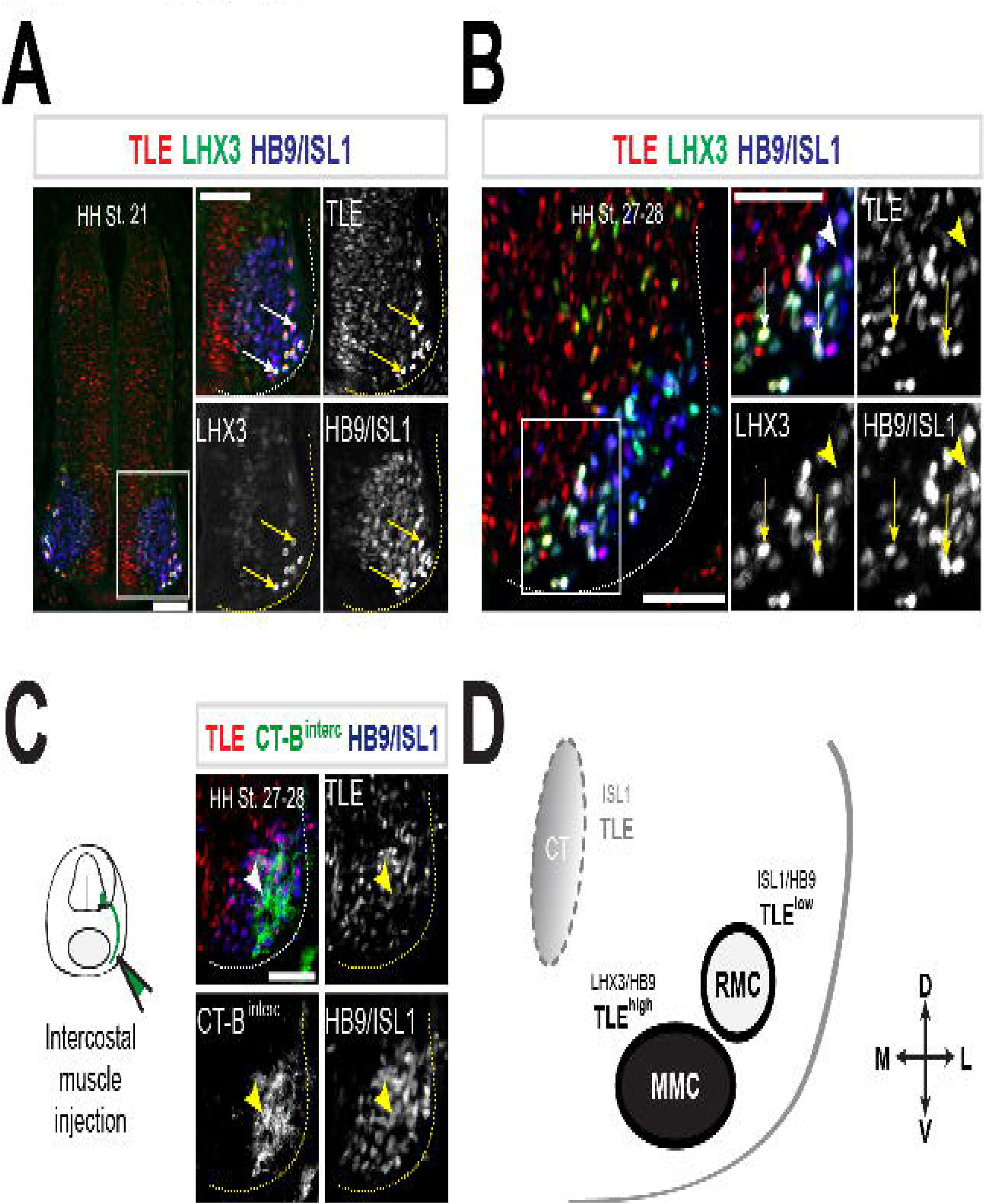
TLE expression in developing thoracic motor neurons. (A and B) TLE expression in the developing chick thoracic spinal cord. Expression of TLE and LHX3 and HB9/ISL1 in the thoracic spinal cord of (A) HH stage 21 and (B) HH stage 27-28 embryos. LHX3-positive cells that do not express HB9/ISL1 likely correspond to V2A interneurons. Arrows point to examples of triple-labelled cells, whereas arrowheads point to examples of MNs showing little or no double labelling. (C) TLE expression in HB9/ISL1+ MNs retrogradely labelled by CT-B injection in intercostal muscles of HH Stage 27-28 chick embryos. (D) Schematic summary of TLE expression in postmitotic MNs at cervical and thoracic levels of the mouse and chick spinal cord. TLE^high^ indicates high level of TLE expression;

A similar expression pattern was observed in the thoracic spinal cord of E12.5 mouse embryos, where TLE^high^ MNs also coexpressed LHX3, while TLE^low^ MNs expressed little or no LHX3 and were retrogradely labelled after injection of CT-B into intercostal muscles (Figure S5A-D). Similar results were obtained when antibodies specific for TLE1 and TLE4 were used (Figure S5E and S5F). An analogous situation was detected in the cervical level, where TLE^high^ expression was observed in cells also expressing LHX3 and RUNX1, which coexpression defines a medial subpopulation of MMC neurons in the cervical spinal cord ^37^ (Figure S5G and S5H). In contrast, little or no TLE expression was detected in lateral LHX3^-^ cells expressing the protein SCIP, previously described as a marker of phrenic MNs of the HMC at cervical level ^3,5^ (Figure S5I). A mostly inverse correlation between TLE and SCIP expression was observed at all rostro-caudal levels of the cervical spinal cord (Figure S5J). Together, these observations provide evidence for an evolutionarily conserved correlation of TLE^high^ expression with MMC and TLE^low^ expression with HMC neurons in both chick and mouse spinal cords (Figure 3D).

### Involvement of TLE in axial as opposed to hypaxial motor neuron topology and target muscle innervation

We further explored the involvement of TLE in MN development by characterizing its effect on thoracic MN body settling positions and axonal projections. Differential topographic positioning of the more medial MMC (LHX3^+^) as opposed to the more lateral HMC (LHX3^-^)somatic MNs (HB9/ISL1^+^) of HH Stage 28 embryos was confirmed through recording of their respective cell body position coordinates on a 2-dimentional matrix (Figure 4A). Using this approach, the digitally reconstructed distribution of *in ovo* electroporated CMV∷TLE1 and CMV∷TLE-DN MNs was determined 96 hpe by recording the coordinates of GFP-expressing MNs on an equivalent canvas subdivided into medial and lateral, as well as dorsal and ventral, compartments. When compared to GFP alone (Figure 4B), expression of CMV∷TLE1 resulted in a significant ventromedial accumulation of electroporated MNs (*p* = 0.009, Hotelling’s *T*^*2*^ test; n=8 embryos) (Figure 4C), suggestive of a preferential MMC topology. Conversely, an increase in electroporated MNs located in more lateral positions, suggestive of a HMC topology, was observed in the presence of CMV∷TLE-DN (*p* = 0.087 relative to control GFP embryos; *p* = 0.00034 relative to TLE1 embryos; Hotelling’s *T*^*2*^ test; n=7 embryos) (Figure 4D). This latter phenotype could be observed as early as 48 hpe (*p* = 0.0085 compared to control GFP embryos; *p* = 0.00044 relative to TLE1 embryos; Hotelling’s *T*^*2*^ test; n=5-6 embryos) (Figure S6).

**FIGURE 4.**
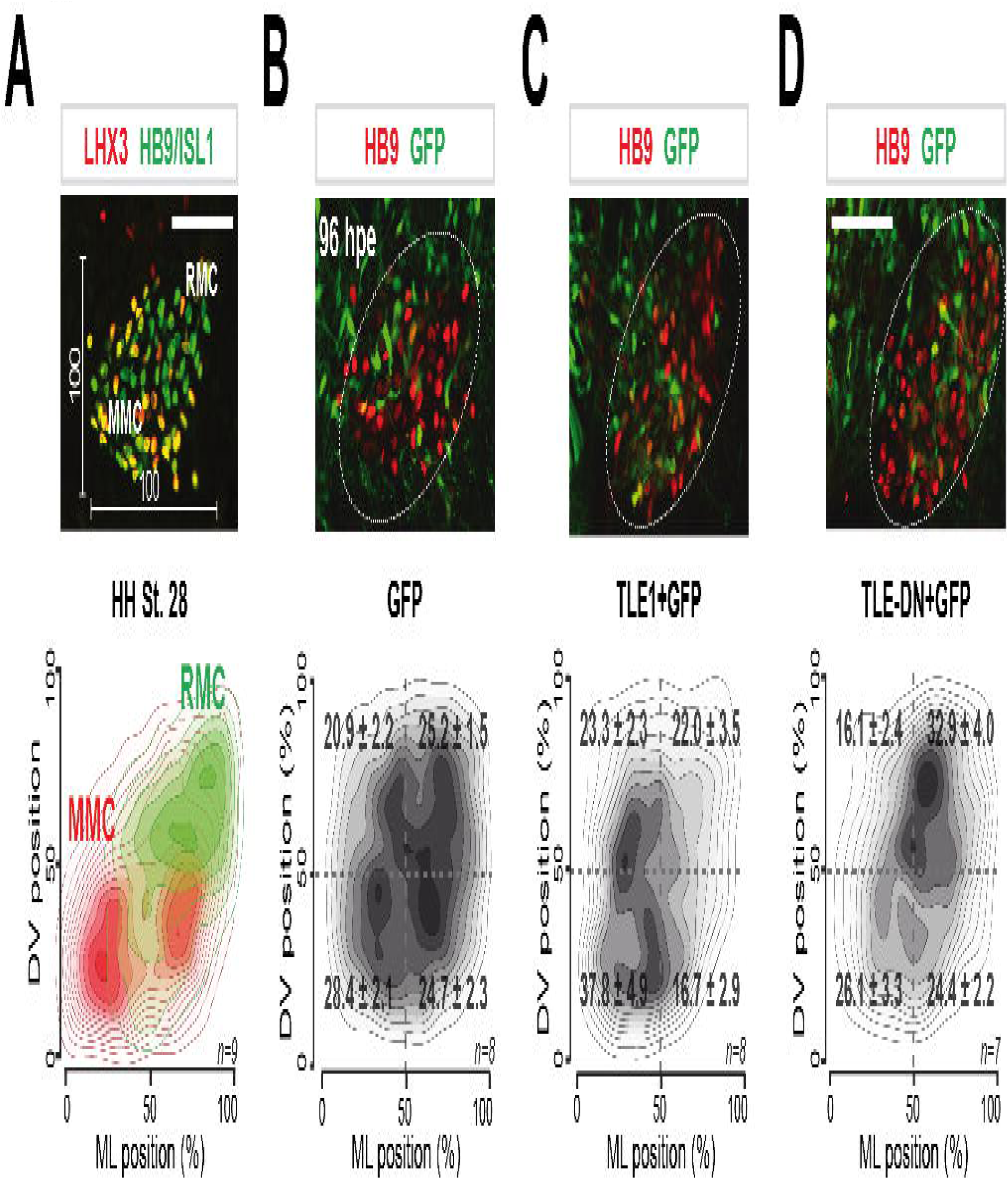
TLE role in somatic motor neuron topology. (A) LHX3 and HB9/ISL1 in representative somatic motor neuron region of a HH Stage 28 chick spinal cords (top). Bottom row: reconstructed mediolateral and dorsoventral distribution of LHX3+ (in red) and LHX3- (in green) somatic MNs at HH Stage 28. A 100×100 virtual template is represented. Here and after, n indicates the number of embryos analysed. (B-D) Topographic distribution of GFP-expressing MNs in the thoracic spinal cord of TLE1- and TLE-DN-electroporated chick embryos. Top row: expression of HB9 and GFP in representative spinal cord hemisections of chick embryos electroporated with GFP alone (B) or together with CMV∷TLE1 (C) or CMV∷TLE-DN (D) and analysed 96 hpe. Somatic MN region is delineated by dotted lines. Bottom row: reconstructed distribution of GFP-positive somatic MNs at 96 hpe on a 100×100 template. The proportion of GFP-positive MNs located in each quadrant along the dorsoventral and mediolateral axes is indicated as mean ± SEM.

To examine whether the observed changes in MN settling positions were associated with axonal innervation alterations, we performed retrograde axonal labelling experiments from intercostal muscles, targets of HMC neurons at thoracic level (Figure 5A-D). CMV∷TLE1 expression resulted in a significant decrease in MNs innervating intercostal muscles (*p* = 0.0085, Student’s *t* test; n=6 embryos) (Figure 5B). Contrariwise, CMV∷TLE-DN expression led to a considerable increase in the innervation of intercostal muscles (*p* = 0.0066 compared to control GFP, Student’s *t* test; n=6 embryos) (Figure 5D). Virtually opposite results were obtained when we examined the effects of these manipulations on the innervation of axial muscles (targets of MMC neurons) (Figure 5E-H). Taken together, these results suggest that TLE is also involved in mechanisms important for the acquisition of thoracic MMC neuronal cell body settling and axial muscle innervation at the expense of HMC topology and target connectivity.

**FIGURE 5.**
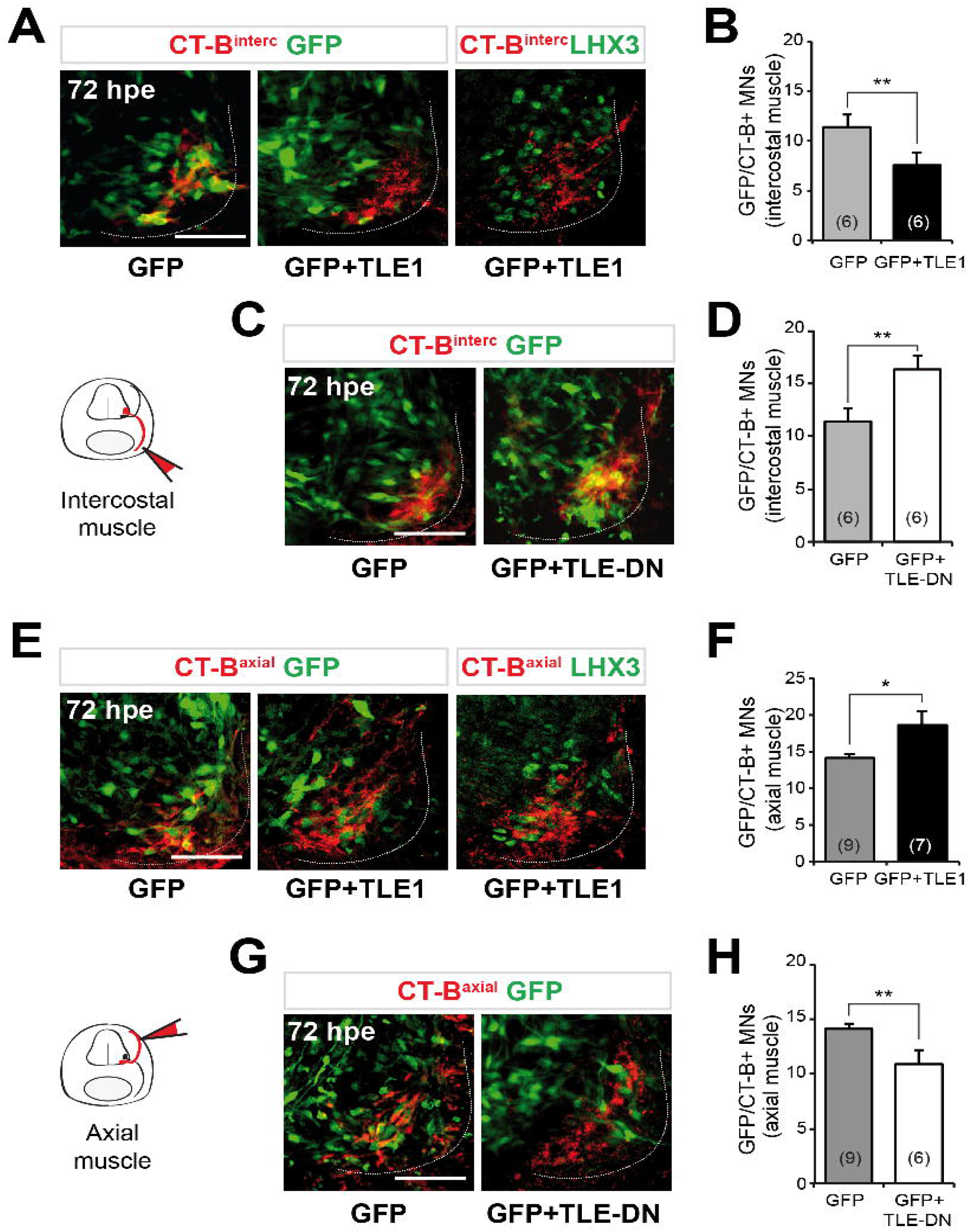
Involvement of TLE in axial and hypaxial motor neuron connectivity. (A-D) LHX3 and GFP expression in chick embryo spinal cords electroporated with GFP alone or together with TLE1 (A) or TLE-DN (C) and then subjected to CT-B retrograde labeling from intercostal muscles at 72 hpe. Quantification of electroporated (GFP+) MNs retrogradely labeled from intercostal muscles with a CT-B tracer in control (GFP), TLE1 (B) and TLE-DN-electroporated embryos (D). Results are shown as mean ± SEM (\**p* < 0.05; \*\**p* < 0.01; n.s., not significant; *t* test). (E-H) CT-B retrograde axonal labeling from axial muscles exogenously expressing GFP alone or together with TLE1 (E) or TLE-DN (G). Quantification of electroporated (GFP+) MNs retrogradely labeled in control (GFP), TLE1 (F) and TLE-DN-electroporated (H) chick embryos spinal cords. Data are mean ± SEM (\**p* < 0.05; \*\**p* < 0.01; n.s., not significant; *t* test).

### Post-mitotic role of TLE in acquisition of axial motor neuron topology and target muscle innervation

Because TLE is expressed in both MN progenitors (Figure 1 and S1) and post-mitotic MNs (Figure 4 and S5), it is conceivable that it might regulate multiple steps of MMC versus HMC development by acting both within and outside the progenitor domain. To explore this possibility, we placed exogenous TLE1 and TLE-DN expression under the control of the neuronal-specific promoter of the *T*α*1* α*-tubulin* (*T*α*1*) gene ^38^ to drive their overexpression selectively in differentiating neurons but not their undifferentiated progenitors. Neither Tα1∷TLE1 nor Tα1∷TLE-DN elicited significant changes in the fractions of MNs displaying MMC or HMC molecular identities, failing to phenocopy the effects of CMV∷TLE1 and CMV∷TLE-DN (Figure S7). In contrast, Tα1∷TLE1 and Tα1∷TLE-DN both impacted on MN three-dimensional distribution and target muscle innervation. Density plot analysis revealed that Tα1∷TLE1 caused a significant accumulation of electroporated HMC neurons in more medial locations (*p* = 0.0000023 when compared to HB9/ISL1^+^/LHX3^-^ neurons in the control non-electroporated sides; *p* = 0.000001 compared to HB9/ISL1^+^/LHX3^-^ neurons electroporated with GFP alone; Hotelling’s *T*^*2*^ test; n=6 embryos) (Figure 6A, 6B and Figure S8). Conversely, we observed that Tα1∷TLE1-DN caused an accumulation of HB9^+^ LHX3^+^ MMC neurons in more lateral locations (*p* = 0.035 compared to MMC MNs in the non-electroporated sides; *p* = 0.014 compared to MMC MNs electroporated with GFP alone; Hotelling’s *T*^*2*^ test; n=5 embryos) (Figure S8). Consistent with these results, Tα1∷TLE1 also impacted on HMC innervation, causing a significant decrease in the number of electroporated MNs labelled by CT-B retrograde transport from intercostal muscles (*p* = 0.0095, Student’s *t* test; n=5 embryos) (Figure 6C and 6D).

**FIGURE 6.**
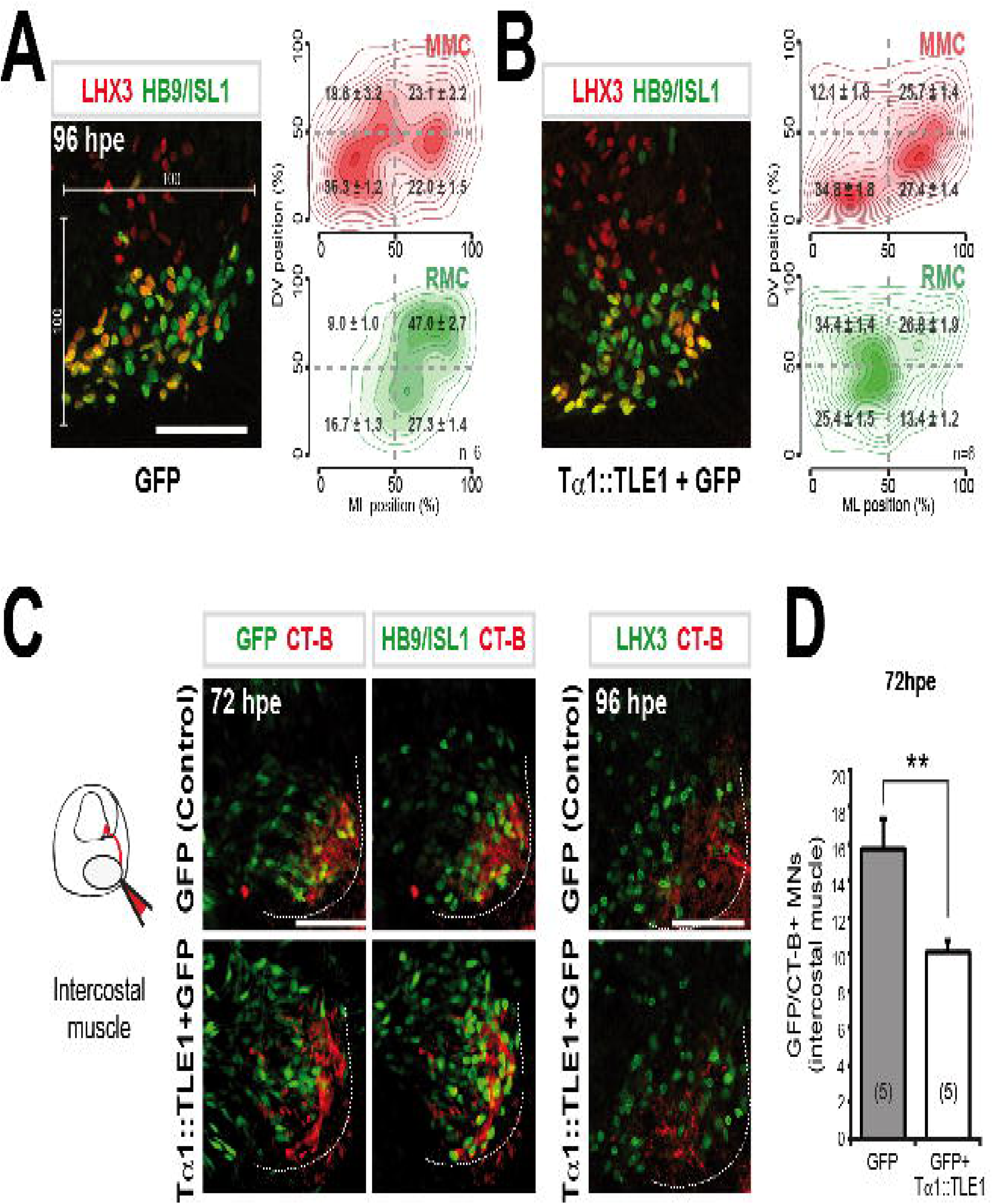
Postmitotic function of TLE in axial and hypaxial motor neuron development. (A and B) Topographic organization of thoracic somatic MNs in chick embryos electroporated with GFP (A) or Tα1∷TLE1 (B). Images on the left show the expression of LHX3 and HB9/ISL1 in representative spinal cord hemisections at 96 hpe. Right columns show digitally reconstructed distributions of somatic MNs on a 100×100 template. The proportion of GFP-positive MNs located in each quadrant along the dorsoventral and mediolateral axes is indicated as mean ± SEM. (C) HB9/ISL1, LHX3, and GFP expression in chick embryo spinal cords exogenously expressing GFP alone or together with Tα1∷TLE1 and subjected to CT-B retrograde axonal labeling from intercostal muscles at 72 or 96 hpe, as indicated. Note the medial displacement of retrogradely labeled RMC neurons in the presence of Tα1∷TLE1. (D) Quantification of electroporated (GFP+) MNs retrogradely labeled with a CT-B tracer following injection into intercostal muscles in control (GFP) or Tα1∷TLE1 electroporated embryos. Control corresponds to proportion in GFP embryos. Results are shown as mean ± SEM (\*\**p* < 0.01; n.s., not significant; *t* test).

Taken together, these findings suggest that TLE acts sequentially during thoracic MN development, first by promoting MMC and repressing HMC neuron subtype specification within the progenitor domain, and then by controlling axial versus hypaxial MN topology and target muscle connectivity choices during post-mitotic development.

## DISCUSSION

In this study, we have accumulated evidence suggesting that the activity of the TLE transcriptional co-repressor is important at multiple steps, both within the progenitor domain and the mantle zone, during the development of postural MMC neurons and respiratory HMC neurons. TLE is first involved in promoting the decision of nascent MNs to acquire a MMC identity, at the expense of the HMC fate. TLE then continues to support the developmental progression toward a MMC fate by specifying the topology and connectivity of MMC neurons whilst suppressing the acquisition of post-mitotic HMC neuron traits. Together with the previous demonstration of an important function of TLE in ventral neural tube patterning ^9,29^, these findings identify TLE as a key regulator of multiple mechanisms underlying MN emergence, subtype specification and axonal innervation choices. The present studies also suggest that TLE is involved in promoting the MMC fate in the thoracic spinal cord, at least in part, by cooperating with non-canonical WNT signalling in MN progenitor cells. This conclusion is based on several observations. Forced exogenous TLE expression phenocopies the generation of supernumerary LHX3^+^ MMC neurons, and concomitant loss of HMC neurons, elicited by overexpression of WNT5A. These effects are observed when exogenous TLE expression is placed under the control of the CMV promoter, which is active in neural progenitors, but not the *T*α*1* promoter, which is active in neuronal cells, indicating a role in progenitor cells. Moreover, dominant inhibition of endogenous TLE causes a reduction of thoracic spinal cord MMC neurons resembling the decrease in MMC neurons elicited by the reduction of *WNT4/5* transcripts ^16^. More importantly, TLE dominant inhibition blocks the ability of exogenous WNT5A to promote MMC, and decrease HMC, diversification, providing evidence that WNT5A acts at least in part in a TLE-dependent manner during this developmental process. Based on these combined observations, we propose that non-canonical WNT signalling acts together with TLE within the pMN domain to integrate extrinsic cues with intrinsic programs of transcriptional regulation resulting in the implementation of thoracic MMC neuronal differentiation programs at the expense of the HMC fate.

The expression of the non-canonical *WNT* genes, *WNT4, WNT5A* and *WNT5B*, defines a ventral ^high^ to dorsal ^low^ gradient in the ventral spinal cord during the acquisition of MN columnar identities ^16^. This situation suggests that the most ventrally located progenitors in the pMN domain are most likely to generate MMC neurons, whereas those progenitors located dorsally, and thus having lower non-canonical WNT activity, are slated to give rise to HMC neurons in the thoracic spinal cord. This model is consistent with the fact that MNs born ectopically from the most ventral part of the neural tube, as is the case in *NKX2.2*-deficient mice, adopt preferentially a MMC fate ^16,39^. In this regard, TLE expression is particularly robust in the most ventral spinal cord, suggesting the existence of a non-canonical WNT^high^ and TLE^high^ condition optimal for ventral MMC fate specification in the absence of NKX2.2. TLE may act to achieve this developmental goal downstream of non-canonical WNT, at least in part, by contributing to the regulation of PAX6 expression. PAX6 is expressed in an almost converse ventral ^low^ to dorsal ^high^ gradient within the pMN domain and TLE is important for correct PAX6 expression pattern in the ventral spinal cord, as shown by the demonstration that dominant-inhibition of endogenous TLE causes a ventral PAX6 expansion, whereas forced TLE overexpression but also WNT5A restricts PAX6 expression more dorsally (Figure S2) ^29^. Thus, it is conceivable that a graded non-canonical WNT:TLE pathway acts, in part, to limit PAX6 expression ventrally in cells fated to become MMC neurons. This possibility is in potential agreement with the demonstration that canonical WNT activates *PAX6* expression in neocortical progenitor cells via β-CATENIN:TCF^34^. Thus, TLE might act to suppress PAX6 in ventral neural tube progenitor cells at least in part through its involvement in the ability of WNT5A to inhibit β-CATENIN:TCF-mediated transactivation. In agreement with this scenario, TLE seems to be involved in modulating competing canonical (dorsal) and non-canonical (ventral) WNT activities involved in establishing a ventral^low^ to dorsal^high^ PAX6 gradient, resulting in PAX6^low^ or PAX6^high^ progenitors giving rise to MMC or HMC neurons, respectively (data not shown). Consistent with these observations, high PAX6 levels play a role in specifying dorsal MN identity and connectivity programs in the developing hindbrain ^40-42^.

It was suggested that conditions of robust non-canonical WNT signalling in the ventral spinal cord might be a characteristic of primitive aquatic vertebrates to ensure that most MNs would acquire a MMC-like phenotype to control undulatory movements ^16^. The present results suggest that a non-canonical WNT:TLE mechanism might have evolved from its functions in undulatory movements regulation found in lower species to a role in the diversification of MMC neurons in birds and mammals. This possibility is in potential agreement with studies in the invertebrate nematode, *C. elegans*, showing that the transcription factor UNC-37, the *C. elegans* ortholog of Groucho/TLE proteins, plays important roles in postmitotic MN development. MNs of the nematode neuraxis are diversified following a binary fate choice in which A-type MNs and their B-type sister cells control backward and forward undulatory locomotion. This developmental mechanism shares many similarities with the MMC/HMC fate choice found in vertebrates where TLE likely converts HMC MN topology and connectivity into those of MMC neurons. Notably, the A/B MN fate choice is guided by a genetic program in which UNC-37 converts B MN traits into those of the A-type, thereby contributing to the regulation of the nematode motility ^43-45^.

In vertebrates, like-minded neurons cluster together in common functionality and connectivity groups to facilitate neural circuit formation. This circuit formation is established in the developing spinal cord when newly born MNs migrate and coalesce to form medial or lateral motor columns. This stereotyped spatial distribution of spinal neurons shapes synaptic connectivity and influences spinal circuit formation. In this regard, we have obtained evidence that TLE activity continues to be important in postmitotic MMC neurons that have emerged from the germinative zone. Specified MMC neurons continue to express robust TLE levels as they settle in the mantle zone and project to axial muscle. In contrast, HMC neurons express noticeably lower TLE levels. Dominant inhibition of endogenous TLE in postmitotic MNs inhibits the acquisition of topology typical of the MMC fate, whereas ectopic TLE^high^ expression leads to decreased HMC topology and concomitant loss of hypaxial muscle innervation. These findings suggest that TLE acts in developing thoracic MMC neurons to prevent cell body settling and innervation choices typical of HMC neurons. They suggest further that TLE plays key roles during several steps governing the separation of the MMC and HMC fates in the developing spinal cord, starting from a role in the specification of MN progenitors cells with separate developmental potential and continuing with roles in the correct topological and muscle innervation choices of the MNs that arise from separate progenitor subdomains. As a result, TLE contributes at different levels to MN generation, topology, presynaptic connectivity to appropriate sensory neuron and interneuron counterparts, and target muscle innervation in the thoracic spinal cord.

In conclusion, these studies have identified TLE as a key regulator of the coordination of MN identity, topology and spinal connectivity programs critical for motor and respiratory functions, implicating this protein family in the establishment of accurate spinal neural networks.

## MATERIALS AND METHODS

### Chick embryo *in ovo* electroporation

Fertilized White Leghorn chicken eggs (Couvoir Simetin Hatchery, Mirabel, QC, Canada) were incubated at 37–38°C in a humidified 1550 Hatcher Incubator (GQF Manufacturing Company, Savannah, GA) until the required developmental stages. *In ovo* electroporation of chick embryos was performed at HH stages 14–16 of embryonic development ^37^. Embryos were injected and electroporated *in ovo* at thoracic neural tube level under a Discovery.V8 stereomicroscope (Zeiss, Toronto, ON, Canada) using the following plasmids: *pCMV2-FLAG-TLE1, pCMV2-FLAG-TLE-DN* (Todd et al, 2012), *pCMV2-FLAG-TLE1(L743F), pCMV2-FLAG-TLE1(R534A*) ^35^, *pcDNA3-PAX6* (#36054) and *pcDNA3-WNT5A-V5* (#35930) (Addgene, Cambridge, MA). All constructs were injected into the chick neural tube together with a *pCAG-GFP* plasmid at a suboptimal concentration ratio of 4:1, as described ^37^. *pT*α*1-FLAG-TLE1* and *pT*α*1-FLAG-TLE-DN* expression plasmids were generated by PCR amplification, followed by subcloning into the *Pst-I* site in a Tα1-promoter-containing plasmid ^38^. Electroporated embryos were allowed to develop to specific stages of development, then dissected, fixed in 4% paraformaldehyde for 30-100 min, and cryoprotected in 30% sucrose for subsequent cryosectioning and immunofluorescence analysis ^46^.

### Transcription assays in *in ovo* electroporated chick embryos

Chick embryos were *in ovo* electroporated at HH stage 14-16 with a TOPFLASH (1 μg/μl) or FOPFLASH (1 μg/μl) luciferase reporter construct ^32^ containing wild-type or mutated TCF/LEF binding sites in the absence or presence of the following DNAs plasmids: *pMyc-TCF3* (0.5 μg/μl), *pCi-neo-*β*-CATENIN* (1 μg/μl), *pCMV2-FLAG-TLE1* (0.5 μg/μl), *pCMV2-FLAG-TLE-DN (*1-2 μg/μl), *pCMV-WNT5A-V5 (*0.5 μg/μl), *pCMV2-FLAG-TLE1(L743F)* (0.5-1 μg/μl), *pCMV2-FLAG-TLE1(R534A)* (1 μg/μl). A GFP-expression plasmid was also co-electroporated. GFP-positive neural tubes were dissected at 18 hpe, homogenized and subjected to determination of luciferase activity as described ^47^. In each case, a *pCMV-*β*gal* plasmid was used to normalize for transfection efficiency. Data were collected from 9-18 dissected embryos per experimental condition.

### Immunofluorescence

Immunofluorescene staining was performed as previously described ^46^. The following primary antibodies were used: rabbit anti-GFP (1:1000; Invitrogen, Carlsbad, CA), rabbit anti-PAX6 (1:500; Covance, Emeryville, CA), rabbit anti-OLIG2 (1:500; Milipore, Billerica, MA), mouse anti-PAX6 (1:10), mouse anti-HB9 (1:10), mouse anti-ISL1 (1:10) (Developmental Studies Hybridoma Bank, Iowa City, IA), rat panTLE (1:10) ^29-31^, rabbit anti-TLE1 (1:500) ^48,49^, rabbit anti-TLE4 (1:500) ^48^, rabbit (1:10,000) and guinea pig (1:5000) anti-SCIP ^3,5^, rabbit anti-RUNX1 (1:1000; Epitomics, Toronto, ON), rabbit anti-TCF4 (1:250; Cell signalling, Beverly, MA), rabbit anti-nNOS (1:1000; ImmunoStar, Hudson, WI). Secondary antibodies included Alexa488 and Alexa555-conjugated anti-mouse or anti-rabbit antibodies (Molecular Probes, Invitrogen, Carlsbad, CA). After single- or double-labelling, sections were mounted with Fluoromount-G (SouthernBiotech, Birmingham, AL). Digital images were captured using Northern Eclipse software (Empix Imaging, Inc., Mississauga, ON, Canada) controlling a Digital Video Camera (DVC, Austin, TX) mounted on an Axioskop 2 or an imager M1 fluorescence microscope (Zeiss, Toronto, ON, Canada). Quantitative analysis was performed using Image J software (National Institutes of Health, Bethesda, MD). Fluorescent signal intensity in Figure 1 and S2 were measured using Image J software as described (http://imagej.net/Image_Intensity_Processing).

### Retrograde axonal labelling

HH stage 27-28 chick embryos were collected into ice-cold PBS, and axial or intercostal muscles were injected with a solution of Cholera Toxin B Subunit-conjugated Alexa555 (Molecular Probes, Life Technologies, Burlington, ON, Canada), following the manufacturer’s recommendation. Embryos were incubated for 4 h at 30 °C in oxygenated DMEM/F12, followed by fixation, embedding in OCT compound, cryosectioning and immunofluorescence staining. A similar approach was used for retrograde labelling experiments in E12.5 mouse embryos as described previously ^5,37^.

### Motor neuron topology analysis

Consecutive cryosections from thoracic spinal cords of *in ovo* electroporated embryos were imaged by immunofluorescence and the regions containing MMC and HMC MNs were analyzed with Image J. Mediolateral and dorsoventral positions of GFP-expressing LHX3-positive (MMC) and LHX3-negative (HMC) somatic MNs were quantitated by assessing the distance from the medioventral MN region corner as the 0.0 biological point ^50^. MN positions were then normalized to a 100×100 plot (at 96 hpe). Positions collected from ≥6 sections per embryo were plotted in “R” using the 2D density function from the “*ggplot2*” package (R Foundation for Statistical Computing, Vienna, Austria, 2005, http://www.r-project.org). Statistical differences between two different conditions were evaluated by Hoteling’s *T*^*2*^ test using the “Hotelling” R package. Distribution of MNs was also assessed using R by counting the percentage of cells that fell within each of 4 quadrants. Distributions are reported as means ± SEM.

### Statistical analysis

The number of stained cells positive for the markers of interest was determined from ≥5 sections per embryo. Quantitative data were expressed as mean ± SEM. Statistical analysis was performed using *R* statistics package. Significance was examined using the unpaired Student’s *t* test or by performing analysis of variance (ANOVA) followed by Tukey or Bonferroni *post-hoc* test, as indicated. Significance was assessed at *p* value < 0.05.

### Animal procedures

All animal procedures were conducted in accordance with the guidelines of the Canadian Council for Animal Care and were approved by the Animal Care Committee of the Montreal Neurological Institute of McGill University.

## Acknowledgments

We thank Dr. A. Kania for insightful comments and suggestions and for critical reading of the manuscript. We thank Dr. J. Dasen for the kind gift of SCIP antibodies and Dr. J.F. Cloutier for microscopy support. We also thank Rita Lo for excellent technical assistance.

## Author contributions

A.S.C. and S.S. designed and co-wrote the manuscript. A.S.C. performed experiments and analyzed data. R.D., J.W.W., M.B., and M.D. performed and analyzed experiments. Y.T. performed experiments. A.S.C established and developed the project. S.S. directed the project.

## REFERENCES

1 Allan, D. W. & Greer, J. J. Development of phrenic motoneuron morphology in the fetal rat. J Comp Neurol 382, 469–479 (1997).

2 Fetcho, J. R. The spinal motor system in early vertebrates and some of its evolutionary changes. Brain Behav Evol 40, 82–97 (1992).

3 Dasen, J. S., De Camilli, A., Wang, B., Tucker, P. W. & Jessell, T. M. Hox repertoires for motor neuron diversity and connectivity gated by a single accessory factor, FoxP1. Cell 134, 304-316, doi:10.1016/j.cell.2008.06.019 (2008).

4 Machado, C. B. et al. Reconstruction of phrenic neuron identity in embryonic stem cell-derived motor neurons. Development 141, 784–794, doi:10.1242/dev.097188 (2014).

5 Philippidou, P., Walsh, C. M., Aubin, J., Jeannotte, L. & Dasen, J. S. Sustained Hox5 gene activity is required for respiratory motor neuron development. Nat Neurosci 15, 1636–1644, doi:10.1038/nn.3242 (2012).

6 Rousso, D. L., Gaber, Z. B., Wellik, D., Morrisey, E. E. & Novitch, B. G. Coordinated actions of the forkhead protein Foxp1 and Hox proteins in the columnar organization of spinal motor neurons. Neuron 59, 226–240, doi:10.1016/j.neuron.2008.06.025 (2008).

7 Nichols, N. L. et al. Ventilatory control in ALS. Respir Physiol Neurobiol 189, 429–437, doi:10.1016/j.resp.2013.05.016 (2013).

8 Briscoe, J., Pierani, A., Jessell, T. M. & Ericson, J. A homeodomain protein code specifies progenitor cell identity and neuronal fate in the ventral neural tube. Cell 101, 435–445 (2000).

9 Muhr, J., Andersson, E., Persson, M., Jessell, T. M. & Ericson, J. Groucho-mediated transcriptional repression establishes progenitor cell pattern and neuronal fate in the ventral neural tube. Cell 104, 861–873 (2001).

10 Novitch, B. G., Chen, A. I. & Jessell, T. M. Coordinate regulation of motor neuron subtype identity and pan-neuronal properties by the bHLH repressor Olig2. Neuron 31, 773–789 (2001).

11 Sander, M. et al. Ventral neural patterning by Nkx homeobox genes: Nkx6.1 controls somatic motor neuron and ventral interneuron fates. Genes Dev 14, 2134–2139 (2000).

12 Shirasaki, R. & Pfaff, S. L. Transcriptional codes and the control of neuronal identity. Annu Rev Neurosci 25, 251–281, doi:10.1146/annurev.neuro.25.112701.142916 (2002).

13 Arber, S. et al. Requirement for the homeobox gene Hb9 in the consolidation of motor neuron identity. Neuron 23, 659–674 (1999).

14 Pfaff, S. L., Mendelsohn, M., Stewart, C. L., Edlund, T. & Jessell, T. M. Requirement for LIM homeobox gene Isl1 in motor neuron generation reveals a motor neuron-dependent step in interneuron differentiation. Cell 84, 309–320 (1996).

15 Thaler, J. P. et al. A postmitotic role for Isl-class LIM homeodomain proteins in the assignment of visceral spinal motor neuron identity. Neuron 41, 337–350 (2004).

16 Agalliu, D., Takada, S., Agalliu, I., McMahon, A. P. & Jessell, T. M. Motor neurons with axial muscle projections specified by Wnt4/5 signaling. Neuron 61, 708–720, doi:10.1016/j.neuron.2008.12.026 (2009).

17 Hollyday, M., McMahon, J. A. & McMahon, A. P. Wnt expression patterns in chick embryo nervous system. Mech Dev 52, 9–25 (1995).

18 Clevers, H. & van de Wetering, M. TCF/LEF factor earn their wings. Trends Genet 13, 485–489 (1997).

19 Mikels, A. J. & Nusse, R. Purified Wnt5a protein activates or inhibits beta-catenin-TCF signaling depending on receptor context. PLoS Biol 4, e115, doi:10.1371/journal.pbio.0040115 (2006).

20 van Amerongen, R., Fuerer, C., Mizutani, M. & Nusse, R. Wnt5a can both activate and repress Wnt/beta-catenin signaling during mouse embryonic development. Dev Biol 369, 101–114, doi:10.1016/j.ydbio.2012.06.020 (2012).

21 Bielen, H. & Houart, C. The Wnt cries many: Wnt regulation of neurogenesis through tissue patterning, proliferation, and asymmetric cell division. Dev Neurobiol 74, 772–780, doi:10.1002/dneu.22168 (2014).

22 Gomez-Orte, E., Saenz-Narciso, B., Moreno, S. & Cabello, J. Multiple functions of the noncanonical Wnt pathway. Trends Genet 29, 545–553, doi:10.1016/j.tig.2013.06.003 (2013).

23 Ishitani, T. et al. The TAK1-NLK mitogen-activated protein kinase cascade functions in the Wnt-5a/Ca(2+) pathway to antagonize Wnt/beta-catenin signaling. Mol Cell Biol 23, 131–139 (2003).

24 Ishitani, T. et al. The TAK1-NLK-MAPK-related pathway antagonizes signalling between beta-catenin and transcription factor TCF. Nature 399, 798–802, doi:10.1038/21674 (1999).

25 Cavallo, R. A. et al. Drosophila Tcf and Groucho interact to repress Wingless signalling activity. Nature 395, 604–608, doi:10.1038/26982 (1998).

26 Chodaparambil, J. V. et al. Molecular functions of the TLE tetramerization domain in Wnt target gene repression. EMBO J 33, 719–731, doi:10.1002/embj.201387188 (2014).

27 Clevers, H. Wnt/beta-catenin signaling in development and disease. Cell 127, 469–480, doi:10.1016/j.cell.2006.10.018 (2006).

28 Nusse, R. WNT targets. Repression and activation. Trends Genet 15, 1–3 (1999).

29 Todd, K. J. et al. Establishment of motor neuron-V3 interneuron progenitor domain boundary in ventral spinal cord requires Groucho-mediated transcriptional corepression. PLoS One 7, e31176, doi:10.1371/journal.pone.0031176 (2012).

30 Stifani, S., Blaumueller, C. M., Redhead, N. J., Hill, R. E. & Artavanis-Tsakonas, S. Human homologs of a Drosophila Enhancer of split gene product define a novel family of nuclear proteins. Nat Genet 2, 343, doi:10.1038/ng1292-343a (1992).

31 Nuthall, H. N., Joachim, K. & Stifani, S. Phosphorylation of serine 239 of Groucho/TLE1 by protein kinase CK2 is important for inhibition of neuronal differentiation. Mol Cell Biol 24, 8395–8407, doi:10.1128/MCB.24.19.8395-8407.2004 (2004).

32 Levanon, D. et al. Transcriptional repression by AML1 and LEF-1 is mediated by the TLE/Groucho corepressors. Proc Natl Acad Sci U S A 95, 11590–11595 (1998).

33 Roose, J. et al. The Xenopus Wnt effector XTcf-3 interacts with Groucho-related transcriptional repressors. Nature 395, 608–612, doi:10.1038/26989 (1998).

34 Gan, Q. et al. Pax6 mediates ss-catenin signaling for self-renewal and neurogenesis by neocortical radial glial stem cells. Stem Cells 32, 45–58, doi:10.1002/stem.1561 (2014).

35 Buscarlet, M., Perin, A., Laing, A., Brickman, J. M. & Stifani, S. Inhibition of cortical neuron differentiation by Groucho/TLE1 requires interaction with WRPW, but not Eh1, repressor peptides. J Biol Chem 283, 24881–24888, doi:10.1074/jbc.M800722200 (2008).

36 Alaynick, W. A., Jessell, T. M. & Pfaff, S. L. SnapShot: spinal cord development. Cell 146, 178–178 e171, doi:10.1016/j.cell.2011.06.038 (2011).

37 Stifani, N. et al. Suppression of interneuron programs and maintenance of selected spinal motor neuron fates by the transcription factor AML1/Runx1. Proc Natl Acad Sci U S A 105, 6451–6456, doi:10.1073/pnas.0711299105 (2008).

38 Gloster, A. et al. The T alpha 1 alpha-tubulin promoter specifies gene expression as a function of neuronal growth and regeneration in transgenic mice. J Neurosci 14, 7319–7330 (1994).

39 Briscoe, J. et al. Homeobox gene Nkx2.2 and specification of neuronal identity by graded Sonic hedgehog signalling. Nature 398, 622–627, doi:10.1038/19315 (1999).

40 Ericson, J. et al. Pax6 controls progenitor cell identity and neuronal fate in response to graded Shh signaling. Cell 90, 169–180 (1997).

41 Osumi, N. et al. Pax-6 is involved in the specification of hindbrain motor neuron subtype. Development 124, 2961–2972 (1997).

42 Takahashi, M. & Osumi, N. Pax6 regulates specification of ventral neurone subtypes in the hindbrain by establishing progenitor domains. Development 129, 1327–1338 (2002).

43 Miller, D. M., 3rd, Niemeyer, C. J. & Chitkara, P. Dominant unc-37 mutations suppress the movement defect of a homeodomain mutation in unc-4, a neural specificity gene in Caenorhabditis elegans. Genetics 135, 741–753 (1993).

44 Pflugrad, A., Meir, J. Y., Barnes, T. M. & Miller, D. M., 3rd. The Groucho-like transcription factor UNC-37 functions with the neural specificity gene unc-4 to govern motor neuron identity in C. elegans. Development 124, 1699–1709 (1997).

45 Winnier, A. R. et al. UNC-4/UNC-37-dependent repression of motor neuron-specific genes controls synaptic choice in Caenorhabditis elegans. Genes Dev 13, 2774–2786 (1999).

46 Methot, L. et al. Interaction and antagonistic roles of NF-kappaB and Hes6 in the regulation of cortical neurogenesis. Mol Cell Biol 33, 2797–2808, doi:10.1128/MCB.01610-12 (2013).

47 Ciarapica, R. et al. Prolyl isomerase Pin1 and protein kinase HIPK2 cooperate to promote cortical neurogenesis by suppressing Groucho/TLE:Hes1-mediated inhibition of neuronal differentiation. Cell Death Differ 21, 321–332, doi:10.1038/cdd.2013.160 (2014).

48 Yao, J. et al. Combinatorial expression patterns of individual TLE proteins during cell determination and differentiation suggest non-redundant functions for mammalian homologs of Drosophila Groucho. Dev Growth Differ 40, 133–146 (1998).

49 Yao, J. et al. Disrupted development of the cerebral hemispheres in transgenic mice expressing the mammalian Groucho homologue transducin-like-enhancer of split 1 in postmitotic neurons. Mech Dev 93, 105–115 (2000).

50 Tripodi, M., Stepien, A. E. & Arber, S. Motor antagonism exposed by spatial segregation and timing of neurogenesis. Nature 479, 61–66, doi:10.1038/nature10538 (2011).

