## Supplemental Figures for "Groucho/TLE opposes axial to hypaxial motor neuron development"

### Figure S1

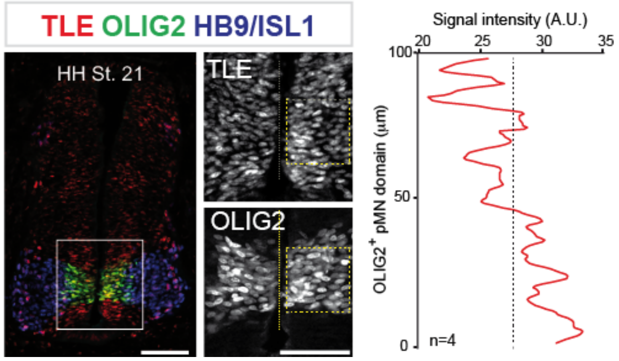


**FIGURE S1. TLE expression in developing thoracic motor neurons**

Expression of TLE, OLIG2 and HB9/ISL1 in HH21 chick spinal cords. Also shown on the right column, fluorescence signal intensity of TLE protein levels within the OLIG2+ pMN progenitor domain of HH Stage 21 chick embryos (n=4 embryos). The *x*-axis depicts signal level of TLE and the *y*-axis the distance (µm) from the bottom limit of the OLIG2 domain.

The boxed areas are shown at higher magnification in columns depicting single channels. Scale bars correspond to 100 µm.

### Figure S2

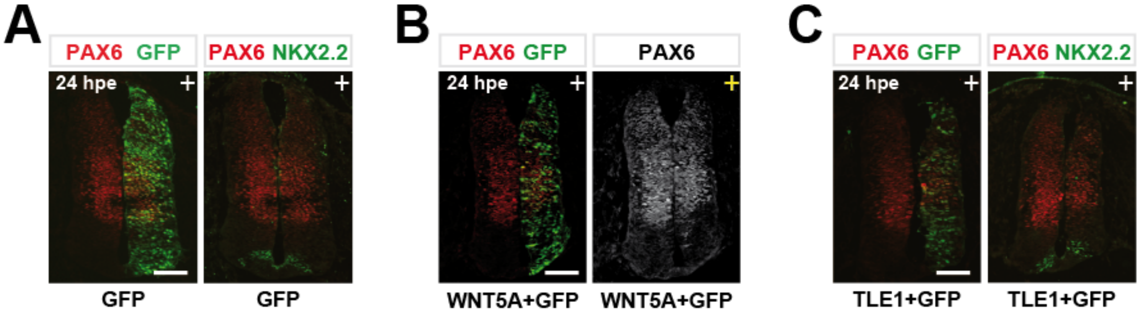


**Figure S2. Exogenous WNT5A and TLE effect on ventral PAX6 expression.**

(A-C) Expression of PAX6, NKX2.2 and GFP in representative sections of chick spinal cords subjected to *in ovo* electroporation with plasmids expressing (A) GFP alone, (B) WNT5A+GFP, or (C) TLE1+GFP and then analysed 24 hpe (HH Stage 21). Here the "+" indicates the electroporated side. Scale bar, 100 µm

### Figure S3

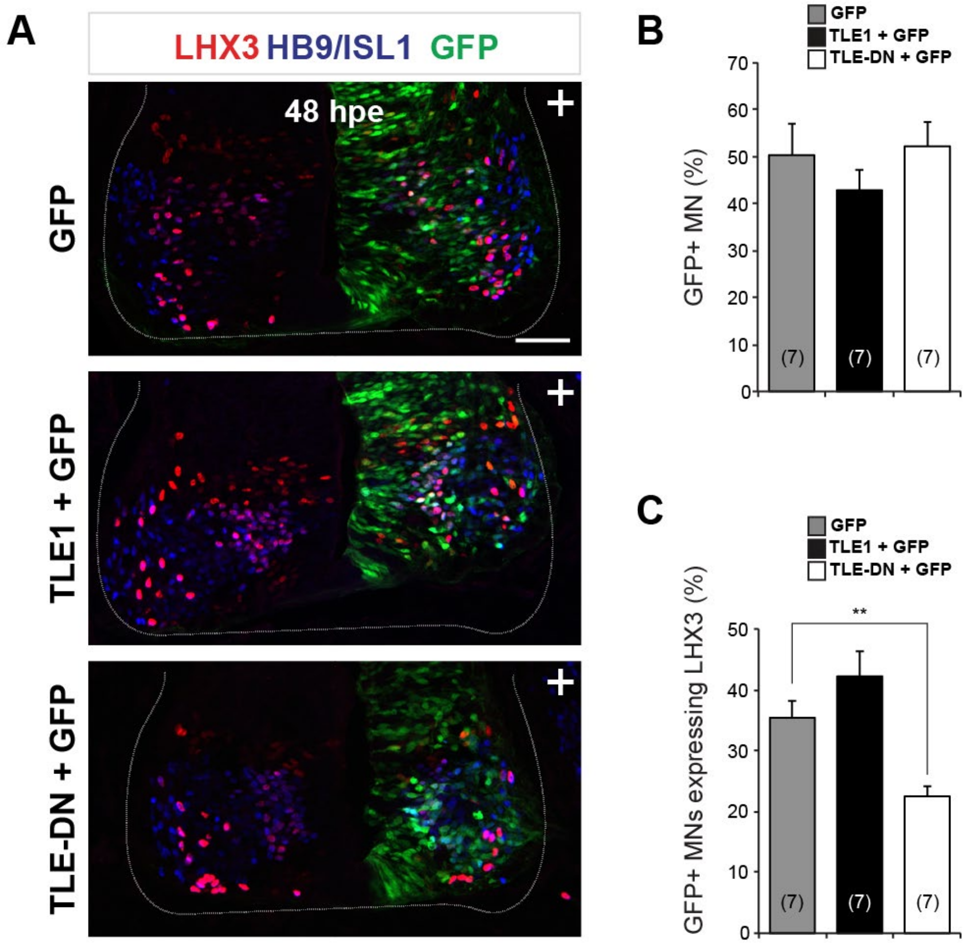


**Figure S3. Role of TLE in thoracic motor neuron subtype specification**

(A) Expression of LHX3, HB9/ISL1 and GFP in representative hemi-sections of chick spinal cords subjected to *in ovo* electroporation with plasmids expressing GFP alone or together with TLE1 or TLE-DN and then analysed 48 hours post-electroporation (HH stage 24). At HH stage 24 both newly born generic MNs and MMC MNs express HB9/ISl1 and LHX3. Scale bar, 100 µm.

(B and C) Quantification of thoracic GFP-posititve (B) total MNs and (C) MNs expressing LHX3 in chick embryo spinal cords electroporated as indicated. Results are shown as mean ± SEM (***p* < 0.01; ANOVA followed by Tukey *post hoc* test).

### Figure S4

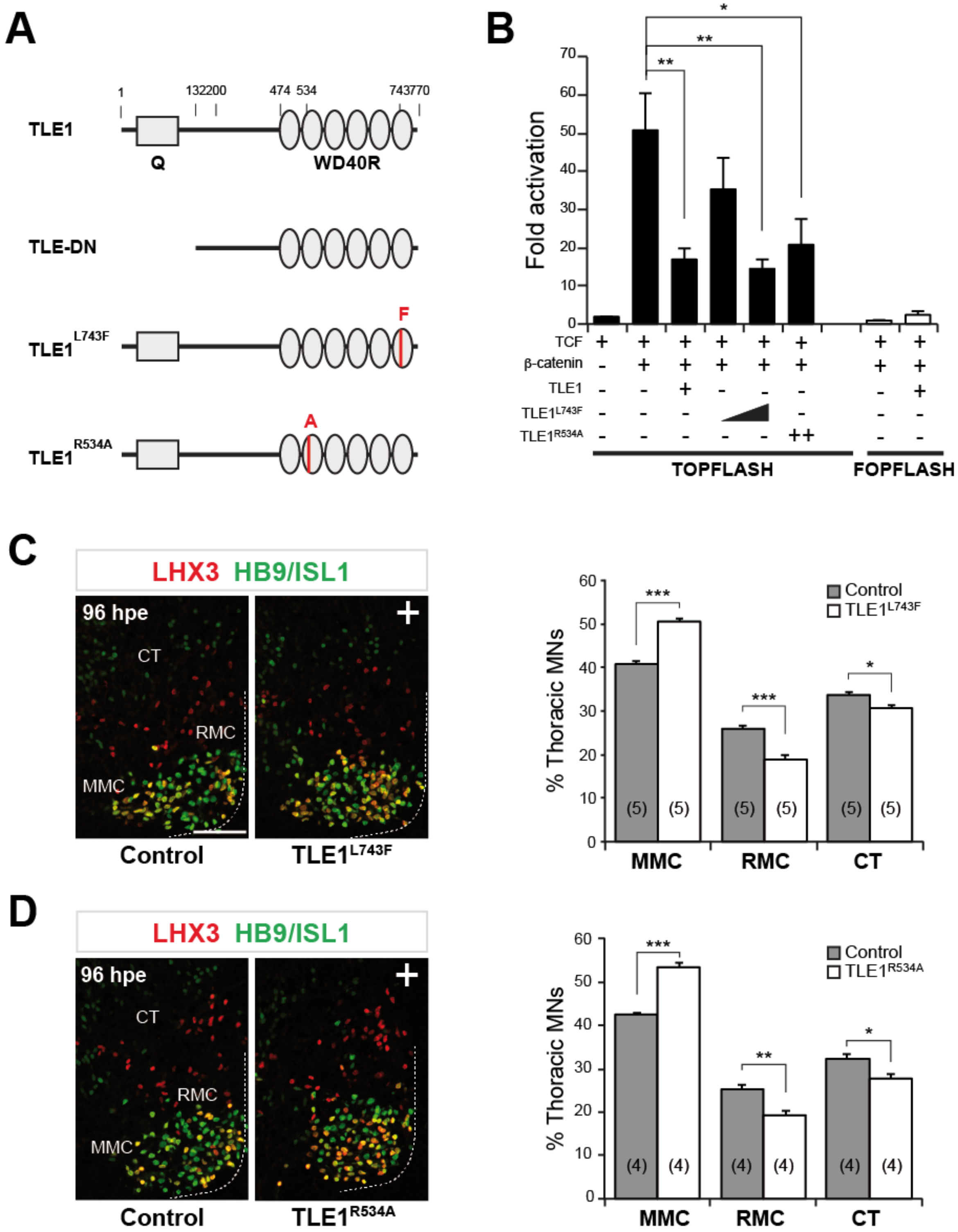
A
B
1
132
200
474
534
743
770
TLE1
Q
WD40R
TLE-DN
F
TLE1L743F
A
TLE1R534A
*
**
**
Fold activation
70
60
50
40
30
20
10
0
TCF
β-catenin
TLE1
TLE1L743F
TLE1R534A
++
TOPFLASH
FOPFLASH
C
LHX3 HB9/ISL1
96 hpe
CT
RMC
MMC
Control
TLE1L743F
% Thoracic MNs
Control
TLE1L743F
***
***
*
(5)
MMC
RMC
CT
D
LHX3 HB9/ISL1
96 hpe
CT
RMC
MMC
Control
TLE1R534A
% Thoracic MNs
Control
TLE1R534A
***
**
*
(4)
MMC
RMC
CT

**Figure S4. TLE WD40 repeat mutations fail to disrupt developing axial and hypaxial balance**

(A) Schematic representation of full-length and mutated TLE proteins used in the present study.

(B) Expression of LHX3 and HB9/ISL1 in representative spinal cord hemi-sections of chick spinal cords subjected to *in ovo* electroporation as indicated and then analysed 96 hours post-electroporation. Scale bar, 100 μm.

(C) Quantification of proportion of thoracic MN columnar subtypes per ventral quadrant of CMV::TLE1(L743F)-electroporated embryos compared to the non-electroporated side (control). Results are shown as mean ± SEM (*$p < 0.05$; ***$p < 0.001$; *t* test).

(D) Expression of LHX3 and HB9/ISL1 in representative spinal cord hemi-sections of chick spinal cords subjected to *in ovo* electroporation as indicated and then analysed 96 hours post-electroporation. Scale bar, 100 μm.

(E) Quantification of proportion of thoracic motor neuron columnar subtypes per ventral quadrant of CMV::TLE1(R534A)-electroporated embryos compared to the non-electroporated side (control). Results are shown as mean ± SEM (*$p < 0.05$; **$p < 0.01$; ***$p < 0.001$).

### Figure S5

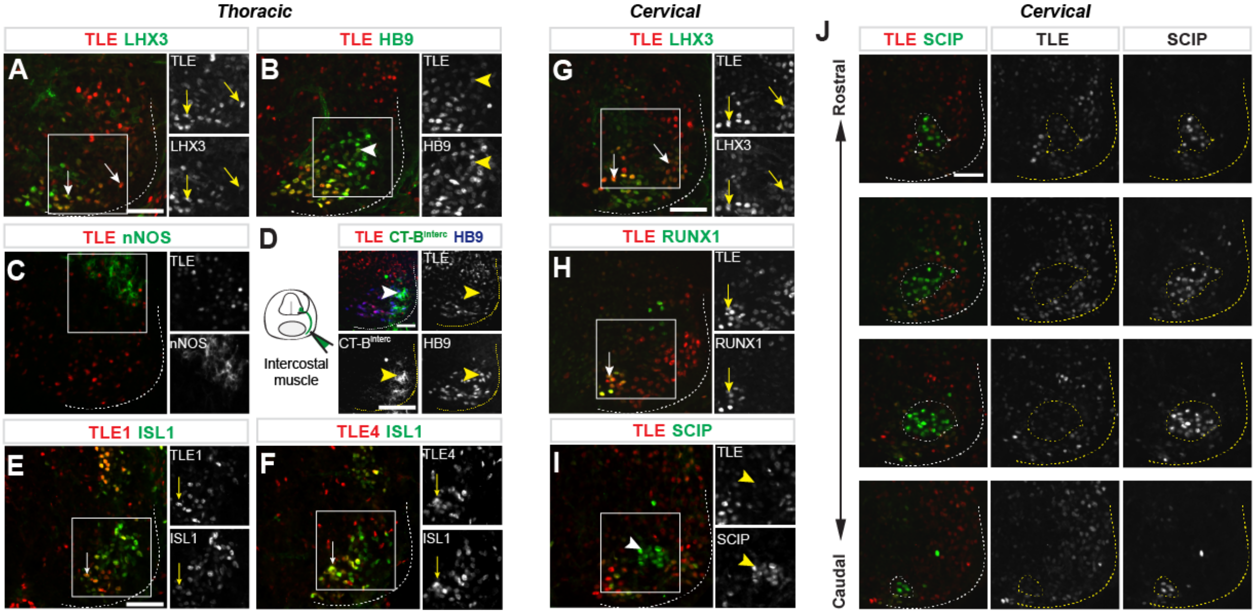

Thoracic
TLE LHX3
A
TLE
LHX3
TLE HB9
B
TLE
HB9
TLE nNOS
C
TLE
nNOS
D
TLE CT-Binterc HB9
TLE
Intercostal muscle
CT-Binterc
HB9
TLE1 ISL1
E
TLE1
ISL1
TLE4 ISL1
F
TLE4
ISL1
Cervical
TLE LHX3
G
TLE
LHX3
TLE RUNX1
H
TLE
RUNX1
TLE SCIP
I
TLE
SCIP
J
Cervical
TLE SCIP
TLE
SCIP
Rostral
Caudal

**Figure S5. TLE expression in the developing thoracic and cervical mouse spinal cord**

(A-C) TLE expression is shown along with LHX3 (A), HB9 (B) and nNOS (C) expression the thoracic spinal cord of developing E12.5 mouse embryos.

(D) TLE expression in intercostal muscles innervating MNs of E12.5 embryos, which were retrogradely labelled by a CT-B tracer

(E and F) Expression of ISL1 and either TLE1 (E) or TLE4 (F) in the thoracic spinal cord of E12.5 mouse embryos.

(G-I) TLE expression in cervical spinal cord of E12.5mouse embryos is shown along with LHX3 (G), RUNX1 (H) and SCIP (I).

(J) Shown are TLE and SCIP expression in serial rostrocaudal sections the cervical spinal cord of E12.5 mouse embryos. Dotted lines delineate regions of high SCIP expression.

Scale bars, 100 μm.

### Figure S6

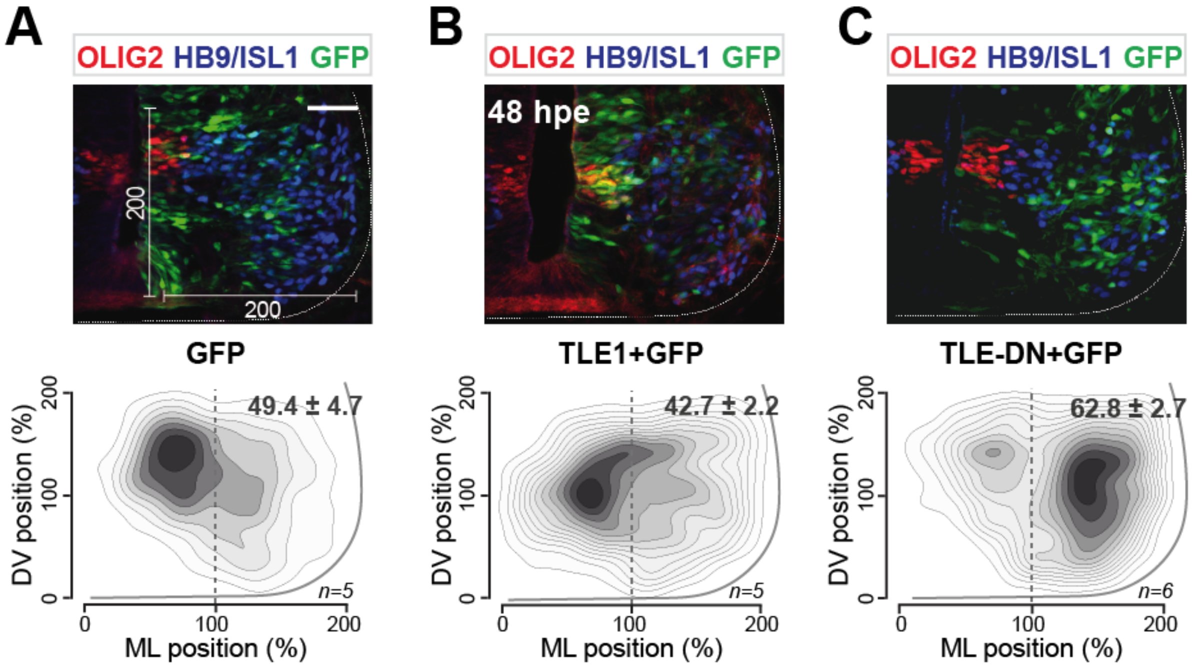


**Figure S6. Topographic distribution of TLE1 and TLE-DN *in ovo* electroporated developing thoracic motor neurons**

(Upper panel) Thoracic MN distribution of GFP-electroporated control embryos (A) as well as TLE1+GFP (C) and TLE-DN+GFP (D) HH24 chick spinal cords. Physical limits of the spinal cord hemisections are delineated by dotted lines. (Lower panel) Topographic distribution of GFP-positive MNs was reconstructed on a 200x200 template. The proportion of GFP+ MNs located in lateral positions of the hemi-sections is indicated.

### Figure S7

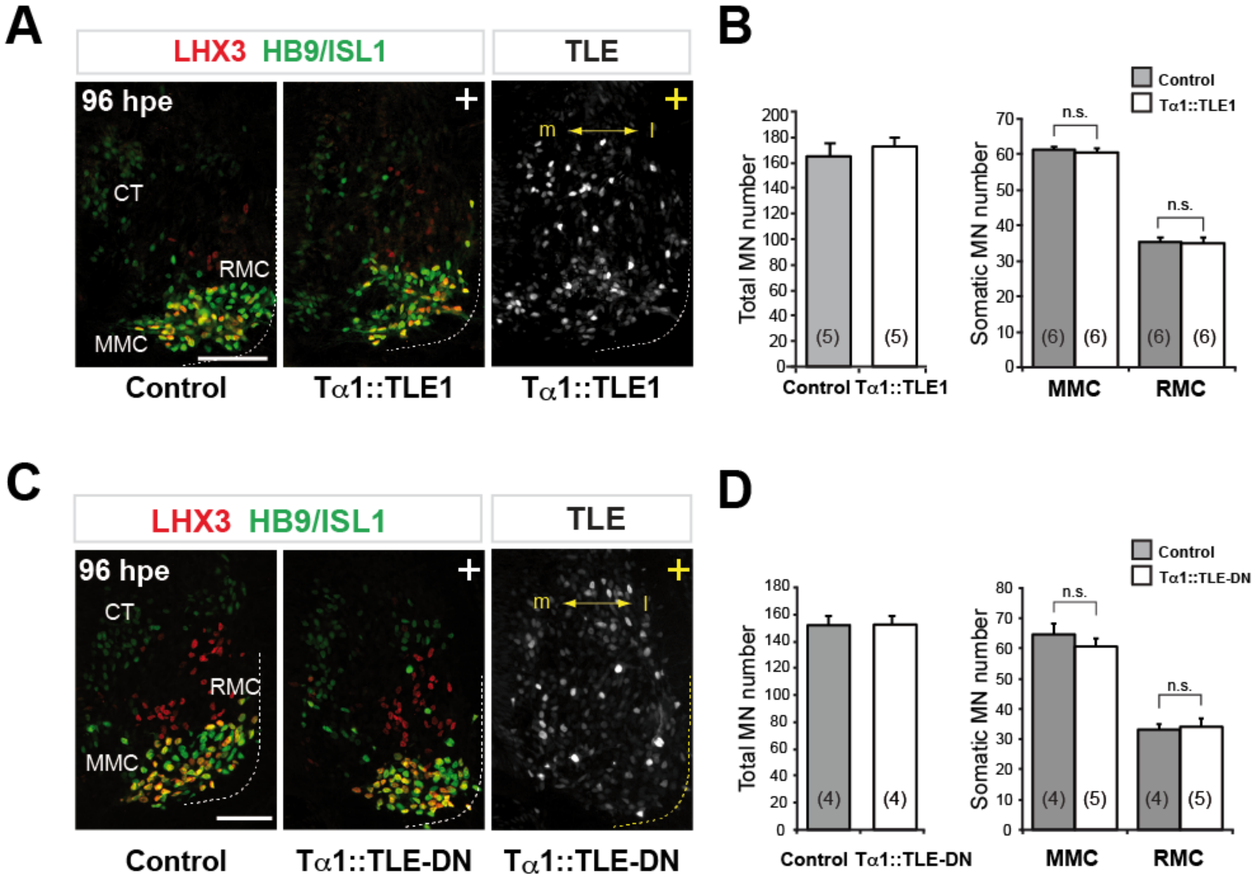


**Figure S7. No TLE post-differentiation effect on thoracic somatic motor neuron identity**

Expression of LHX3, HB9/ISL1 and TLE in representative hemi-sections of chick spinal cords subjected to *in ovo* electroporation in Tα1::TLE1 (A) and Tα1::TLE-DN (C) embryos and then analysed 96 hours post-electroporation.

Quantification of total thoracic MN numbers or somatic MN columnar subtypes as indicated (B and D) at 96 hpe. Control corresponds to the proportion of MNs observed on the non-electroporated side. Results are shown as mean ± SEM (n.s. not significant; *t* test).

Scale bars, 100 μm.

### Figure S8

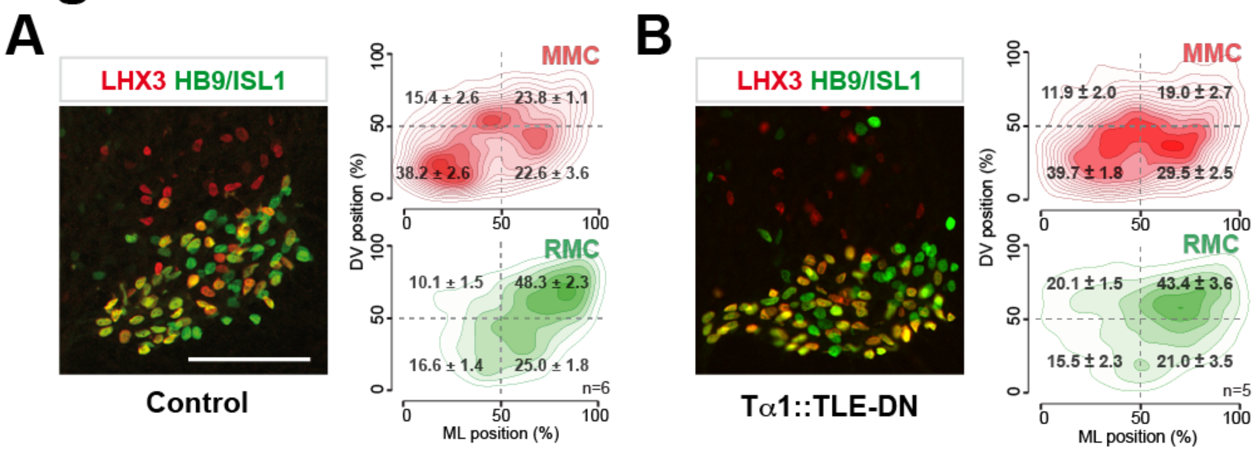


**Figure S8. Postmitotic TLE in axial and hypaxial motor neuron topology**

In both panels, left columns show the expression of LHX3 and HB9/ISL1 to depict the topographic organization of thoracic somatic MNs in representative spinal cord hemisections of non-electroporated chick embryos (A) or embryos electroporated with GFP together with Tα1::TLE-DN (B) analysed 96 hours post-electroporation. Right columns show digitally reconstructed distributions of somatic MNs on a 100x100 template. The proportion of GFP-positive MNs located in each quadrant is indicated as mean ± SEM.
